# Influence of aversive cue detection sensitivity on extinction in adult male rats

**DOI:** 10.1101/2024.04.30.591853

**Authors:** Emma N Cahill, Emily R Sherman, Joseph Jollans, Serena Deiana, Bastian Hengerer

**Affiliations:** Department of Physiology, Development and Neuroscience, University of Cambridge, CB2 3EB, UK; CNS Discovery Research, Boehringer Ingelheim Pharma GmbH & Co. KG, Biberach an der Riss, 88397, Germany; School of Physiology, Pharmacology and Neuroscience, University of Bristol, BS8 1TD, UK

**Keywords:** Fear, Anxiety, Bed nucleus of the stria terminalis, Memory, Extinction, Vigilance, Ultrasonic Vocalization

## Abstract

Threat detection prompts reactions classified either as fear (obvious, predictable, immediate threat) or anxiety (ambiguous, sustained, distant threat). Hypervigilance is a state of sensitivity to threatening stimuli and an attentional bias symptomatic of anxiety disorders. In rodents, threat detection can be measured by freezing behaviour and production of ultrasonic vocalisation (USV) alarm calls. The amygdala is classically associated with fear-like responses, whereas the bed nucleus of the stria terminalis (BNST) has been proposed to be preferentially recruited by anxiogenic stimuli. The conditioned responses triggered by aversive cues can be extinguished through repeated exposure of a subject to the threat stimulus but without any aversive reinforcement. The extent of extinction acquisition and consolidation are notedly variable across individuals. It has been reported that NMDA-type glutamate receptor co-agonists, like D-cycloserine, can enhance extinction consolidation. In the experiments herein, the salience of a threat cue was modified to compare the relative activation of the brain vigilance networks to an obvious cue, and to test whether sensitivity to the aversive cue at such a ‘vigilance screen’ might predict subsequent ability to extinguish conditioned responses. We demonstrated activation of the BNST by a low salience aversive cue. Rats that had the propensity to make alarm ultrasonic vocalisation calls reacted more strongly to aversive cues and had deficits in conditioned freezing extinction. Finally, we demonstrated the potential to enhance extinction consolidation by targeting glycine transmission. Taken together these results demonstrate how threat detection and responses are sensitive to cue salience and can be manipulated by combined pharmacological and behavioural interventions.

**HIGHLIGHTS:** -Auditory cue at low salience revealed attentional bias unrelated to maze behaviour

-Low salience cue recruited activation of the BNST

-Alarm call vocaliser rats had deficit in extinction consolidation

-GlyT1 inhibition enhanced extinction consolidation

## INTRODUCTION

Rodent Pavlovian conditioning tasks are widely used to associate defensive behaviours to otherwise neutral cues for insight into conserved mechanisms of threat detection and memory. Although simplistic, the aversive associations that can be measured as changes in behaviour provide useful models for learning, attention, memory or emotion (1). Hypervigilance is a symptom of anxiety disorders defined by heightened detection and reactivity to potential threat. Hypervigilance is distinct from avoidance, and in some cases could be the source of an avoidance behaviour. Startle responses, induced by a sequence of aversive stimuli provide a means to study how altered state (by a stressful acute stimulus) can lead to exaggerated responses relative to a baseline state. In aversive associative conditioning tasks, variability has been recognised in the levels of responding at test and also during subsequent extinction of those responses (2). Studies have even suggested that such variability may not be only a state dependent effect, but could be a heritable phenotype (3). Extinction strength can be modified by pharmacological and/or behavioural interventions (4,5). Given the requirement of NMDA-type glutamate receptor (NMDAR) signalling for neuronal plasticity, studies explored manipulation of NMDAR to influence the extinction of conditioned defensive responses (6). The NMDAR activity is regulated by the binding of glutamate and the tonic influence of the co-agonists (7). The co-agonist D-cycloserine (DCS) was demonstrated to enhance the consolidation of conditioned freezing extinction in rats (8). The proposed mechanism of action was that NMDAR signalling and the subsequent process of cellular consolidation of an inhibitory memory was enhanced via neuronal plasticity. The capacity for translation to Humans as a pharmacological adjunct to exposure therapy was explored for phobias, PTSD and other anxiety-related disorders but unfortunately, this was met with varying levels of success (9).

We hypothesized that reactivity to a threat cue that was low in relative salience might reflect an attentional bias akin to hypervigilance. We predicted that such a low-salient cue might preferentially activate the circuitry argued to be recruited more by anxiogenic, rather than fear, stimuli. It is argued that more ambiguous or distant threats engages circuitry prominently featuring the bed nuclei of the stria terminalis rather than acute or predicted threats, which engage predominantly the amygdalae nuclei (10), although it is debated how much this reflects a robust difference between anxiety and fear (11,12). We also hypothesized that extinction of conditioned responses could be enhanced by targeting the alternative NMDAR co-agonist to serine, namely glycine, by increasing its availability using a glycine transporter (GlyT1) inhibitor (Bitopertin). Bitopertin dose dependently increases glycine availability in rats (13), and reached phase III clinical trials as a potential Schizophrenia treatment (14). Unfortunately, there was no additional benefit of Bitopertin over placebo in those patient trials, yet no concerns for safety were raised.

These experiments revealed that through modification of aversive conditioning protocols, to develop a vigilance screen test, we demonstrated a recruitment of the BNST by low-salient aversive cues. Furthermore, rats that vocalised during extinction were found to reach typical extinction acquisition levels, but displayed deficits in extinction consolidation and persistence. Finally, systemic treatment with Bitopertin enhanced the consolidation of extinction of threat responses.

## MATERIALS AND METHODS

### Subjects

Subjects were 136 adult male Lister-Hooded rats (Charles River and Envigo, UK) weighing approx. 250–300g at the start of experiments. All animals were housed in groups of four per cage and kept under a reversed 12 h light/dark cycle (lights off 07:00 h until 19:00 h) and were provided with food and water ad libitum, except for during behavioural procedures, which were conducted during the rats’ dark cycle. The rats were randomly assigned to each group and experimenters were blind to group allocation for subsequent analysis. This research was conducted on Project Licence 70/7548 and has been regulated under the Animals (Scientific Procedures) Act 1986 Amendment Regulations 2012 following ethical review by the University of Cambridge Animal Welfare and Ethical Review Body (AWERB).

### Behavioural procedures

For auditory conditioning, four identical operant boxes (29.5 x 32.5 x 23.5cm, MedAssociates) were used. Each with a Plexiglass rear wall, hinged door and roof, with aluminium side walls. The boxes also contained a house light (2.5W, 24V), a speaker (3kHz tone, Sonalert Module with volume control, MedAssociates), ultrasonic microphones (Metris, Netherlands) and a CCTV video camera (Spy Camera model CMOSNC76). The boxes were positioned within sound attenuating chambers and each also contained a ventilation fan, which provided background noise (approx. 60dB). At time zero the house light was turned on. The same box was used for each rat throughout the experiments. The unconditioned stimulus was a mild foot-shock (0.5mA, 0.5s). All training and test sessions were video recorded for off-line behavioural analysis. The percentage of time freezing (absence of movement except for breathing) during 1 min before (Pre CS) and during the CS was manually scored from the videos by two observers.

### Elevated Plus Maze

The elevated plus maze (EPM) was carried out as described previously (18). In brief, rats were tested for exploration of the open arms for a period of 5 mins each. Behaviour was filmed and monitored from an adjacent but separated room. The EPM was carried out seven days before the aversive conditioning procedures started.

### Auditory Conditioning

On Day 0 and 1 rats were habituated to the context for 1 hour. On Day 2, after a baseline period of 20 min three pairings of the 1 min tone (3kHz, 75dB) coterminous with the footshock were presented with 5 min inter-stimulus intervals. At the ‘vigilance screen’, Day 3, after 5 min of baseline the tone was presented for 1 min twice with a 5 min interval, and then the rats remained in the context for a further 5 mins. The volume of the tone was adjusted at the vigilance screen to 75dB for the ‘normal volume’ groups and to 65dB for the ‘low volume’ groups, and recorded using a decibel meter. For the Extinction sessions, after a 2 min period of baseline there were 20 or 10 presentations of the 1 min tone (3kHz, 75dB) depending on the experiment, with 1 min intervals and then rats remained in the context for a further minute. The test of Extinction consolidation was performed 24h later, and also served as a session to test reacquisition of conditioned freezing. After 5 mins of baseline the tone was presented for 1 min (3kHz, 75dB) and a single co-terminus footshock was presented (0.5mA, 0.5s). The rat remained in the context for a further 5 mins. The test of Reacquisition was performed 24hs later, and after a baseline of 5 mins there were two presentations of the 1min tone (3kHz, 75dB) with a 5 min interval and the rat remained in the context for a further 5 mins.

### USV Analysis

An Ultrasound Microphone (Metris, Netherlands) was placed through a hole in the middle of the operant box roof about 30 cm above the shock floor. The microphones were sensitive to frequencies of 15–125 kHz. Vocalization was recorded and analysed using the Sonotrack software (Metris, Netherlands). Call detection was provided by an automatic threshold-based algorithm. Experimenters manually screened the calls automatically detected and classified based on the mean frequency as 22kHz alarm calls or >55kHz calls or cage noise. The number of USV calls and total calling time are reported for the 22kHz calls only.

### Tissue preparation

1.5h after behavioural testing, the rats were injected with sodium pentobarbital (15) and subsequently perfused transcardially with a phosphate buffer solution (PBS) for 2 mins, followed by normal buffered formalin (NBF). The brains were stored in NBF for 48 hours before transfer to a cryoprotective 30% sucrose solution for at least 48hs and then stored at-80’C until sectioning. 30µm sections of the NAc, the BNST and the BLA were then collected using a cryostat (Leica, CM3050S, Lecia Biosystems, Germany). The level of slices to be taken was determined by comparing the gross neuroanatomy with the Rat Brain Atlas (16), with the boundaries of the regions of interest as shown in Figure 2b.

**Figure 1:**
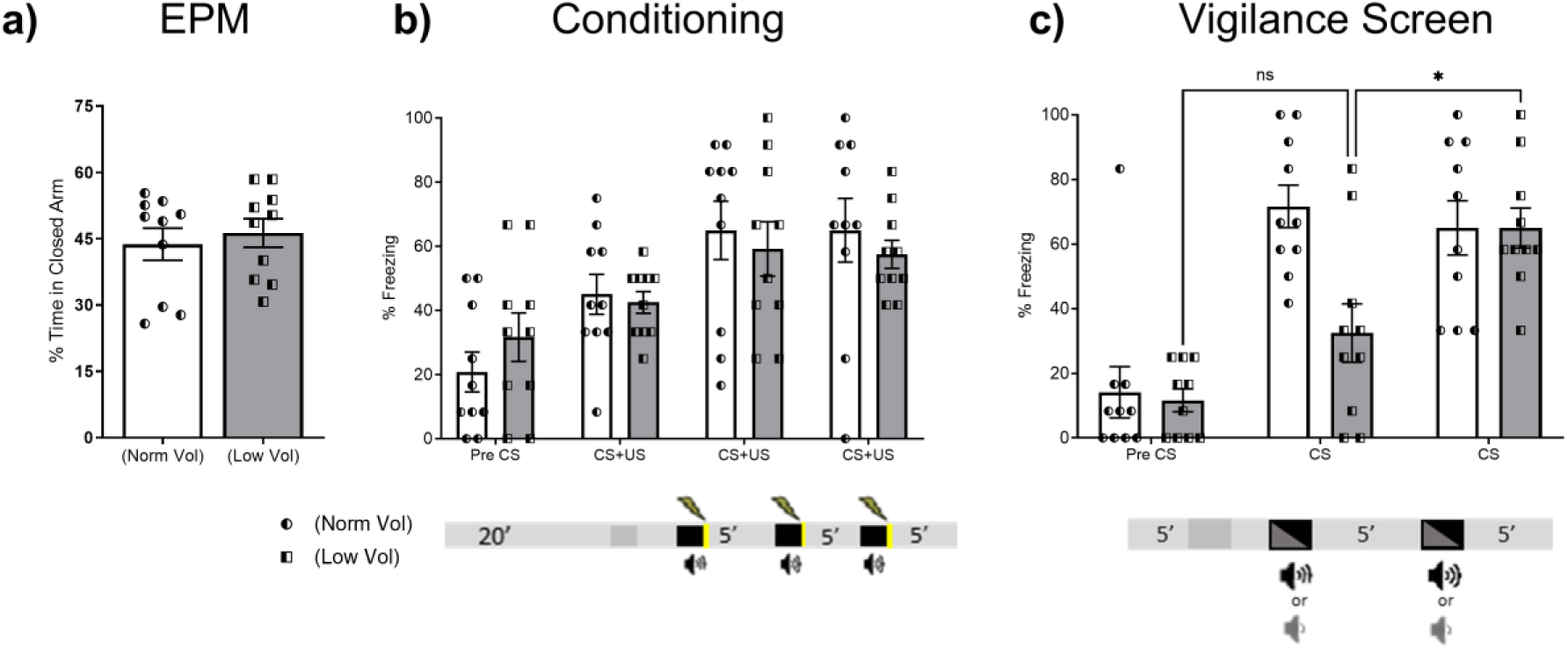
a) Time spent in the closed arm of the EPM for the prospective treatment groups. B) Percentage time of conditioned freezing during aversive conditioning with normal volume tone. Session timings shown in schema. C) Test of reactivity to a low or normal volume tone. Session timings shown in schema. Bars are means +/-SEM, each data point is a rat. *P<0.05.

**Figure 2:**
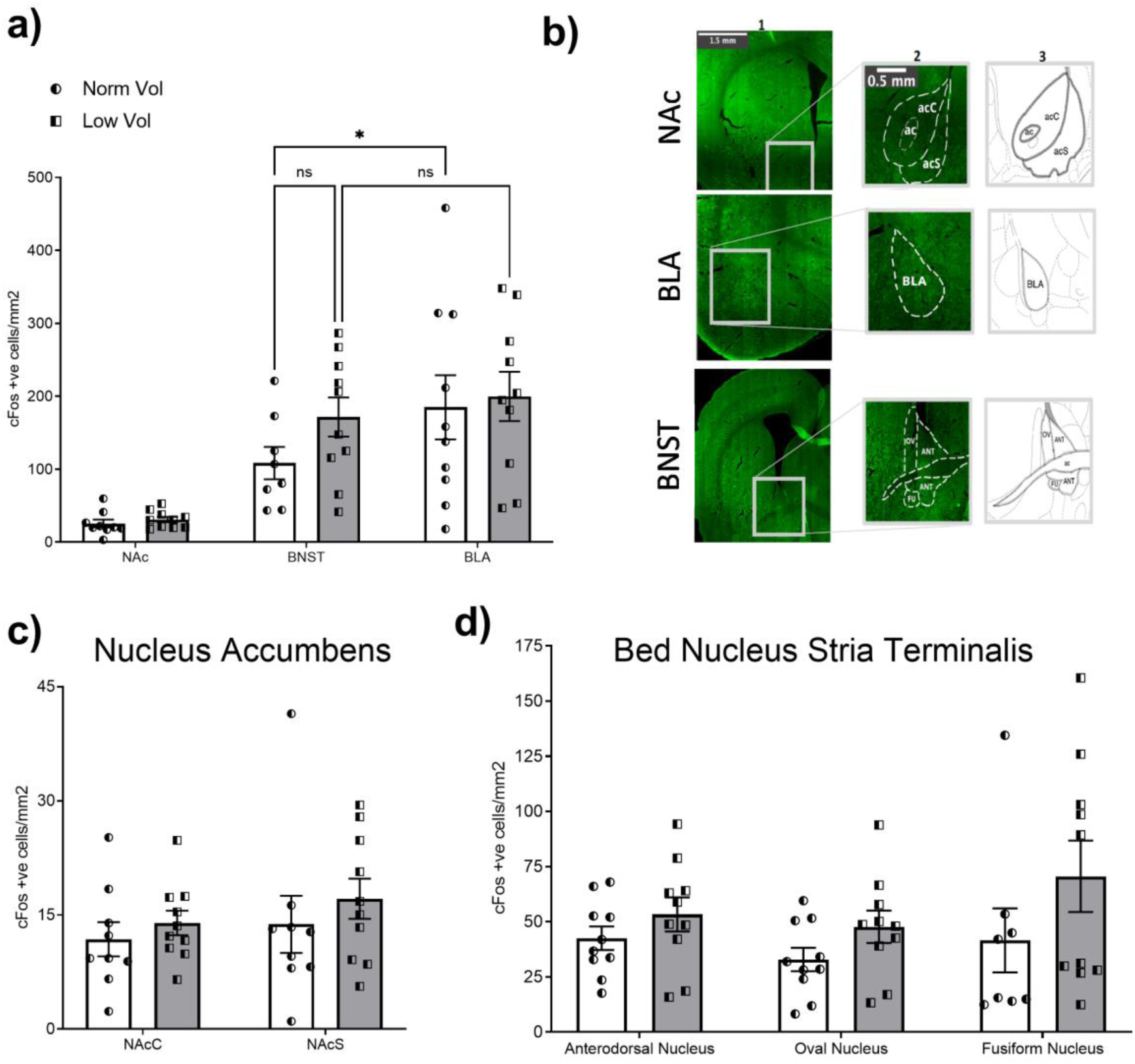
a) Number of cFos positive cells across regions of interest. B) Illustrative images of region of interest boundaries. C) Number of cFos positive cells in the subregions of the NAc. D) Number of cFos positive cells in the subregions of the BNST. Bars are means +/-SEM, each data point is a rat. *P<0.05.

### Immunohistochemistry

Tissue slices were washed 3 times in PBS for 10 mins and then placed in blocking solution (3% normal goat serum, 0.3% triton in 0.1M PBS) for 2 hours at room temperature. Slices were incubated in the primary c-Fos antibody (ABE457, Millipore) for 24 hours at 4’C on a rocker. Following a further 3 washes in PBS, the tissue was incubated in the secondary antibody of goat anti-rabbit IgG Alexa Fluor 488 (ab150077, Abcam, UK) for 2 hours at room temperature. The tissue was washed again three times in PBS and mounted for imaging with medium containing DAPI (ab104139, Abcam). The number of c-Fos positive nuclei in each ROI was quantified using ImageJ software. Tissue damage in the region of interest caused loss of data points for two subjects in the Fusiform nucleus and one subject in the Nucleus Accumbens. Quantifications were calculated as density of c-Fos positive cells per mm2. The value for each individual rat was averaged across hemispheres.

### Drugs and Pharmacokinetics

D-4-amino-3-isoxazolidone (DCS, C3909 Sigma-Aldrich, UK) was prepared in sterile saline solution (6). Bitopertin (S8219, Stratech, UK) was prepared in a 40% 2-hydroxypropyl-β-cyclodextrin in saline solution (17), this vehicle solution was used for the control vehicle comparison groups. Both drugs were administered by intraperitoneal (i.p.) injection, DCS at 1 ml/kg and vehicle or Bitopertin at 2 ml/kg. To analyse plasma availability of drugs post injection, trunk samples of blood were collected from experimentally naïve rats 0.5h after i.p. drug administration. Samples were collected in tubes coated with EDTA and centrifuged. These samples were frozen until subsequent analysis. Plasma and concentrations of Bitopertin and DCS were determined by liquid chromatography coupled to tandem mass spectrometry (LC-MS/MS).

### Statistical Analysis

Data are presented as mean ± SEM, unless otherwise stated. Statistical analyses included t-tests, repeated-measures ANOVA, and planned Tukey’s (across all means) or Šídák’s (independent) comparisons for more than three groups were made using GraphPad Prism (GraphPad Software Inc., La Jolla, CA, USA, version 10.0.2.232). Where Mauchly’s Test of Sphericity indicated the assumption of sphericity had been violated, degrees of freedom were Greenhouse-Geisser corrected. For comparison of categorical data, i.e. the proportion of rats that produced 22kHz vocalisation or not, a Fisher’s Exact Test was performed. The significance level was set at p<0.05. Graphs and figures were generated in GraphPad Prism.

## RESULTS

### An auditory cue at low salience reveals attentional bias for threat detection

Prospective groups were tested in the elevated plus maze (EPM) to characterize initial behaviour in an established task routinely used to measure anxiety-like behaviour (i.e. avoidance of bright space, Fig.1a). No significant difference in time spent in the closed arm was found between the prospective groups (t_18_=0.5165, P=0.6118). Subsequently, both prospective groups increased freezing behaviour to a similar extent over the presentations of the normal volume tone CS and US pairings during conditioning (F_cs(2.261,40.70)_= 16.85, P<0.001, Fig1b). The following day, a vigilance screen was performed where one group were presented with the tone CS twice at a low volume (Low Vol) to reduce its salience. The levels of time spent in the closed arm during the EPM test did not significantly correlate with behaviour at the later Vigilance Screen (P=0.9870), suggesting the two tests measure distinct reactions to threats (18). At the first presentation of the Low Vol CS, freezing responses were significantly lower as a group than those presented a Norm Vol CS as expected (P=0.0083, Fig1c). At the second presentation freezing increased to match Norm Vol levels (F_vol*CS_ (_2,36_) = 7.065, P<0.0026), with both groups demonstrating increased freezing relative to PreCS levels.

### An auditory cue at low salience recruits activation of the BNST

1.5h after the vigilance screen tissue was collected to perform immunohistochemistry. The number of c-Fos positive cells per region of interest (ROI) was highest for the BLA, relative to the BNST and NAc (F_ROI(1.521, 25.10)_ = 26.84, P<0.0001, Fig2a). An overall effect of CS volume across all regions did not reach significance (F_vol_(_1,18_) = 0.7283 P=0.4053). The NAc was examined under the assumption that the area that would not be significantly recruited by threat cues at the vigilance screen, and as expected no subregion difference were noted across the NAcC nor NAcS between the groups (F_vol_(_1,17_) = 0.7283 P=0.4053, Fig2c). Interestingly the c-Fos density in the Low Vol group was as quantitatively high in the BNST as in the BLA, suggesting that a low-salience threat cue activated the BNST as strongly as the BLA. The more obvious, Norm Vol, cue preferentially recruited the BLA over the BNST (P=0.0449), whereas this was not different in the Low Vol group (P=0.5180). When the subregions of the BNST were considered for the influence of cue volume salience overall it reached statistical significance (F_vol_(_1,52_) = 4.906, P=0.0312, Fig2c), but the difference between Norm and Low vol groups within each region did not withstand the planned *a priori* comparison (_Anterodorsal_ P=0.8279, _Oval Nucleus_ P=0.6465, _Fusiform Nucleus_ =0.1563). It appeared that in the most lateral structure, i.e. the Fusiform nuclei, the c-Fos levels significantly correlated negatively with freezing behaviour (n=20, r^2^=0.2598, P=0.0307), which may have drove the overall difference. Taken together, an obvious cue preferentially recruited the BLA over the BNST, but the less salient cue recruited activation of both the BNST and also BLA.

### The production of 22kHz USVs calls reveals hypervigilant responders with deficits in consolidation of extinction

In another cohort of rats the vigilance screen was repeated, and followed by extinction training and a test of reacquisition of the CS-US association to examine whether conditioned freezing during a low salience cue might predict subsequent responding. Both prospective groups conditioned to comparable extents (F_cs(2.879, 86.36) =_ 18.54 P<0.0001, F_group_(_1,30_) <1, Fig3b). Again the Low Volume cue condition lead to significantly lower group levels of freezing but with considerable variability in reaction strength (F_vol*CS_(_2,60_) =3.545, P=0.0351, Fig 3c). The next day, the cue was presented at the normal volume unreinforced to both groups twenty times for acquisition of extinction (Fig 3d). Both groups extinguished freezing over the session to a comparable extent (F_cs(9.244, 277.3)_ = 7.978, P<0.0001). There was a significant interaction between groups and CS number (F_vol*cs_(_19, 570_) = 1.955, P=0.0091) likely driven by the higher responses of the Low Vol group to the first few CS presentations, nonetheless these were not statistically significantly different from the Norm Vol group. The next day, a reacquisition session was performed with the time before the shock in presence of the CS serving as a test of extinction consolidation (Fig3e). The Low Vol group displayed significantly higher freezing behaviour to the cue relative to pre-CS levels (P=0.0002), however overall both groups responded similarly to the CS (F_vol_ (_1, 30_) <1). The next day, the CS was presented twice to examine reacquisition (Fig3f). Both groups reacquired a significant difference between the pre-CS and CS levels of freezing (F_cs (1.645, 49.34)_ = 17.89,P<0.0001). The levels of freezing detected in the vigilance screen by the second CS presentation in the Low Vol group did not significantly correlate with the freezing at the Extinction test (P=0.7255) nor Reacquisition test (P=0.7511). At the Vigilance screen, USV data for four rats was lost due to equipment failure (n=1 from Norm Vol, n=3 Low Vol). Nonetheless, the rats producing alarm calls in the vigilance screen displayed significantly higher levels of freezing (F_Vocalisation_ (_1, 24_) = 10.23, P=0.0039, Fig S1). We therefore looked during the extinction session whether alarm vocalisation might predict heightened reactivity to the CS cue at the reacquisition sessions (Fig4). Indeed, vocal rats were the most reactive in terms of freezing to the CS presentations (F_vocalisation_ (_1, 30_) = 9.177, P=0.0050, Fig4b) with significantly higher levels of freezing at the start of the session (until CS6, F_vocalisation*CS_ (_19, 570_) = 3.270, P<0.0001,), yet they still acquired extinction by the CS20 presentation similar to the levels of the non-vocal rats (Fig4c). Remarkably, the following day the vocal rats persisted in heightened responses to the CS cue relative to the non-vocal rats (F_vocalisation*CS_ (_1, 30_) = 4.581, P=0.0406, Fig4d). The reacquisition of freezing was also significantly higher in the rats that had vocalised during extinction acquisition (F_vocalisation*CS_ (_1, 30_)_=_ 8.835, P=0.0058, Fig4e).

**Figure 3:**
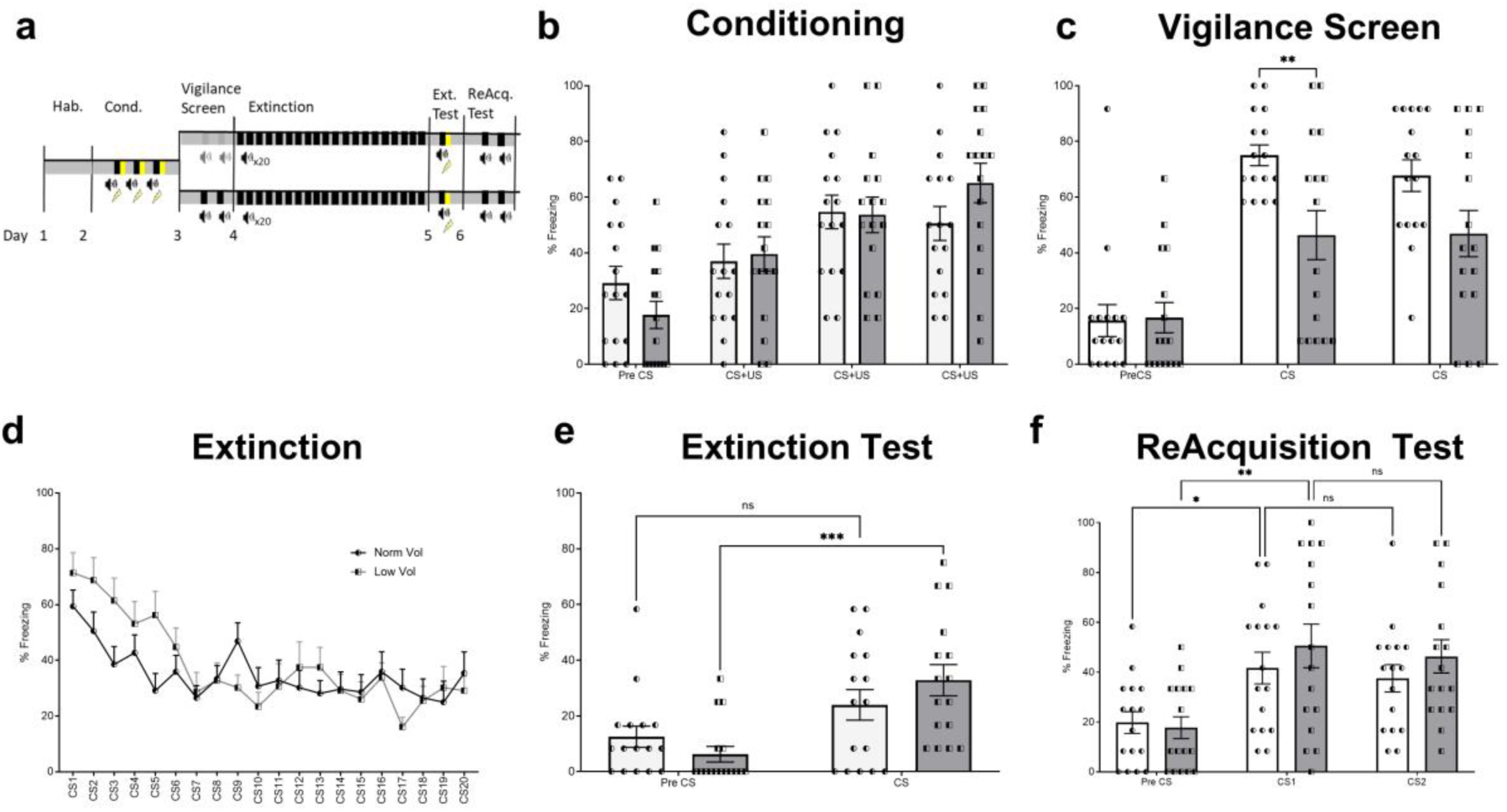
a) Schema of experimental timeline. B) Percentage time of conditioned freezing during aversive conditioning with normal volume tone. C) Test of reactivity to a low or normal volume tone. D) Extinction acquisition across 20 unreinforced normal volume CS presentations. E) Conditioned freezing during the test of Extinction. F) Conditioned freezing after repairing of the CS+US to test Reacquisition. Bars are means +/-SEM, each data point is a rat. *P<0.05.

**Figure 4:**
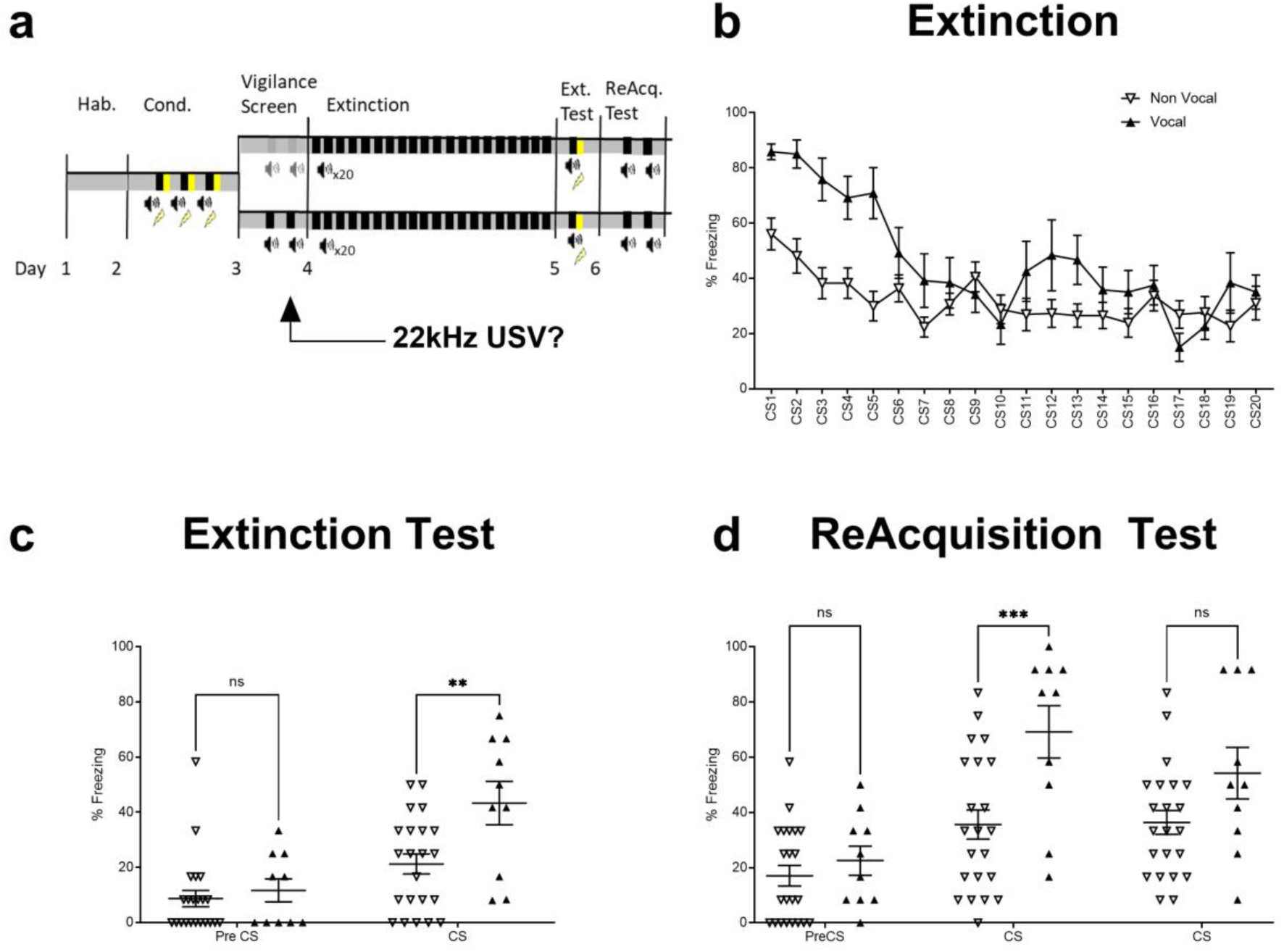
a) Schema of experimental timeline, the propensity to produce 22kHz alarm calls was assessed during the extinction acquisition session. B) Percentage time of conditioned freezing during acquisition of extinction across 20 unreinforced normal volume CS presentations. C) Conditioned freezing during the test of Extinction. D) Conditioned freezing after repairing of the CS+US to test Reacquisition. Bars are means +/-SEM, each data point is a rat. **P<0.001, ***P<0.0001.

### Consolidation of extinction was enhanced by GlyT-1 inhibition

A cohort of experimentally naïve rats were injected with drugs predicted to enhance activation of the glycine co-agonist site of NMDAR to determine plasma availability. One group was injected with D-cycloserine (D-4-amino-3-isoxazolidone; DCS) at a dose demonstrated to enhance extinction (15mg/kg, (19,20)), and the other groups with a dose range of Bitopertin predicted to reach levels to enhance glycine availability (0.6mg, 2mg, 6mg/kg). Trunk blood was collected 0.5h later, and analysis revealed the doses reached predicted increased concentrations in plasma that were associated with raised levels of CSF glycine (Figure 5, (17)). To examine the behavioural effects, DCS and the two higher Bitopertin doses (2mg and 6mg) were systemically administered to rats 0.5h before the beginning of a partial extinction acquisition session (Fig 6a). All prospective groups conditioned to a similar extent (F_cs_ (_3, 180_) = 201.2, P<0.0001, F_drug_ (_3,60_) <1). The proportion of rats that were vocal during extinction acquisition was significantly impacted by drug treatment (Fishers exact test, P=0.0313), which was driven by the high dose of Bitopertin inhibiting vocalisation. There was no significant drug effect on the acquisition of extinction as indicated by conditioned freezing, as all groups decreased to a similar level across the 10 CS presentations (F_cs (4.029, 241.8)_ = 50.76, P<0.0001). The following day however there was significant drug effect on CS-triggered freezing behaviour (F_cs*drug_ (_3, 60_) = 2.887, P=0.0429). The response of DCS-treated group was not significantly different from the vehicle control in contrast with our predictions (P=0.6292), however the 2mg/kg dose of Bitopertin had a significant effect on conditioned freezing (P=0.0352). Whilst there was also a lower level of freezing in the 6mg/kg group, it was not significantly different from the control group (P=0.2087). At the test of reacquisition, there was no significant effect of drug that persisted across the CS presentations (F_drug (3, 56)_ = 0.3065, P=0.8205).

**Figure 5:**
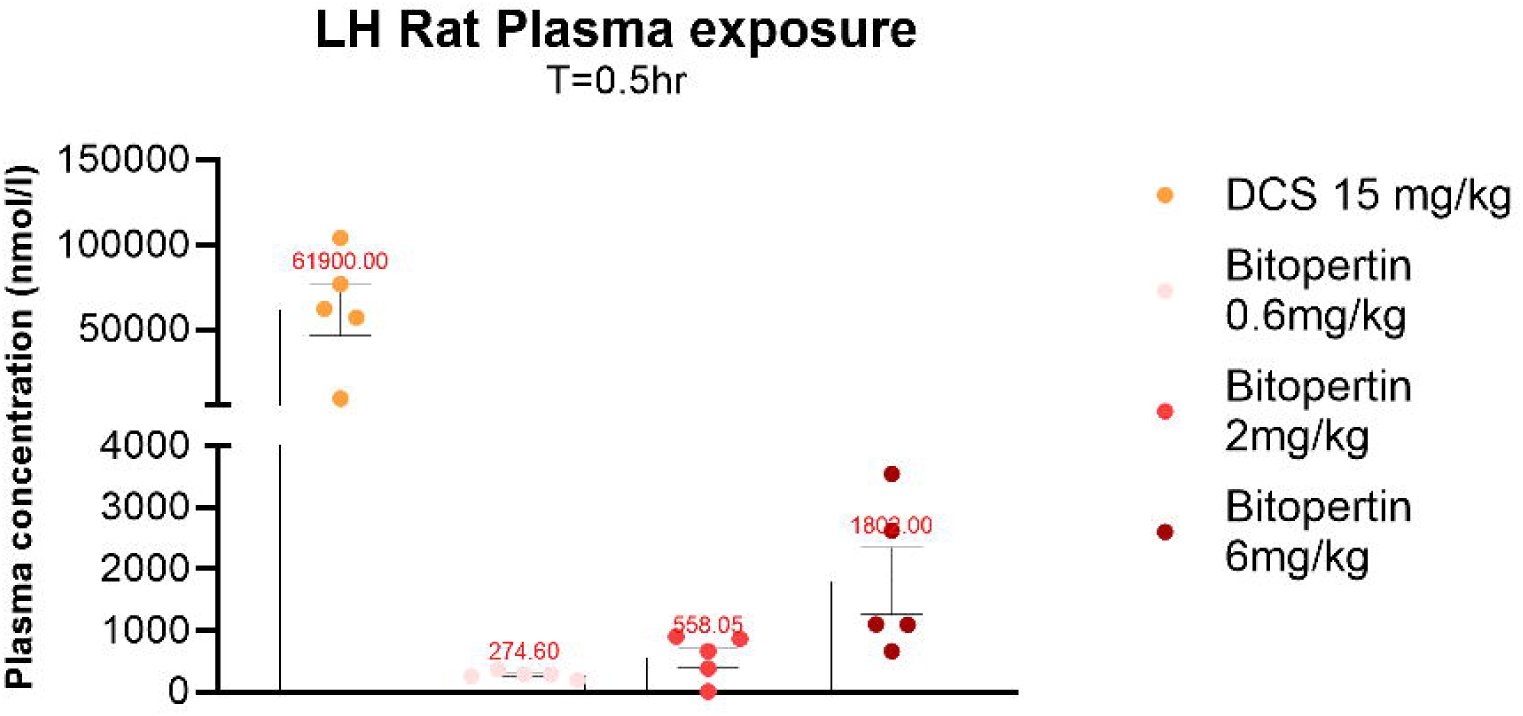
Plasma concentration of drug 0.5h after i.p. administration. Bars are means +/-SEM, each data point is a rat.

**Figure 6:**
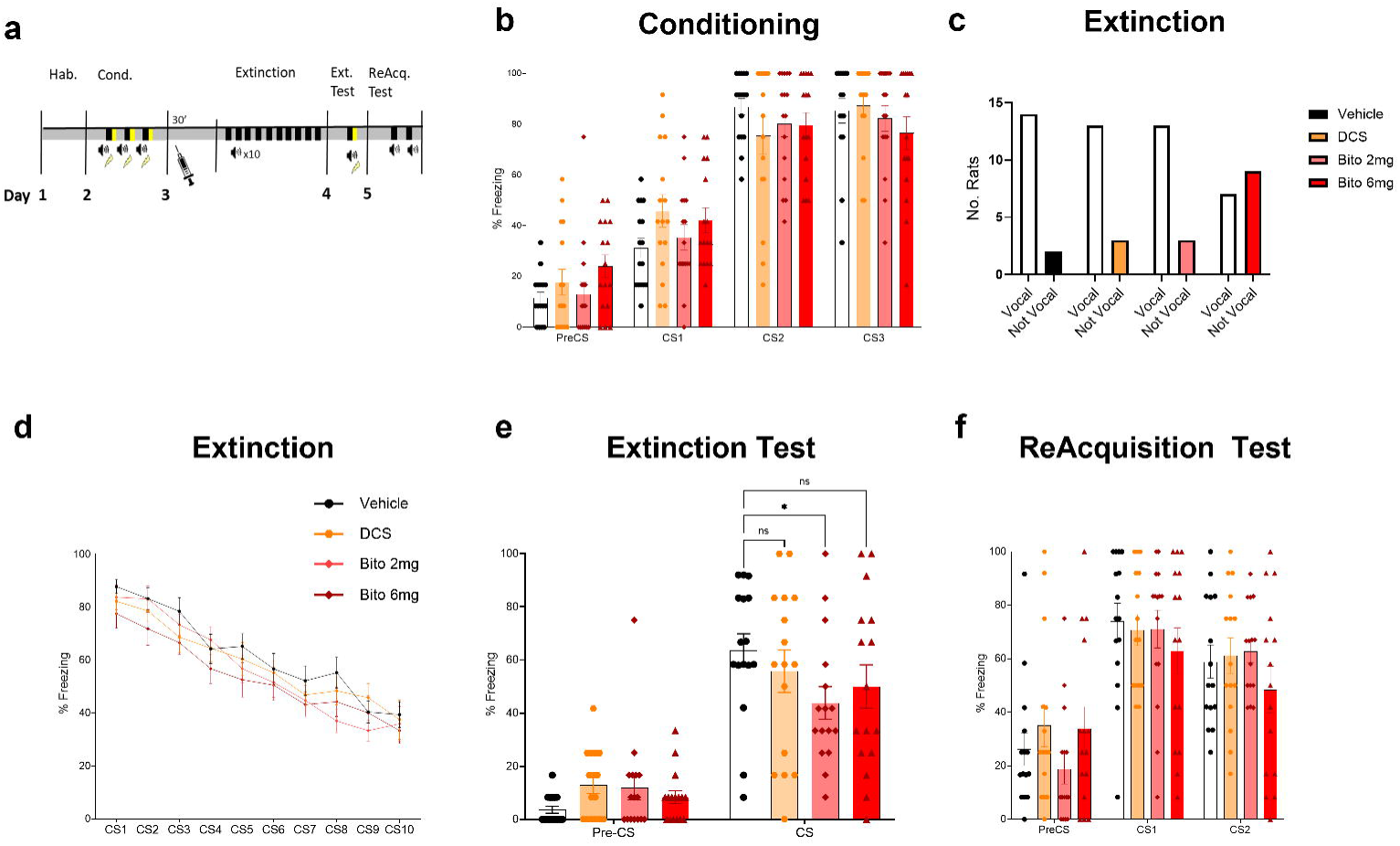
a) Schema of experimental timeline, all cues presented at normal volume. B) Percentage time of conditioned freezing during aversive conditioning. C) Relative proportion of rats that produced 22kHz alarm calls during the acquisition of extinction session after drug treatment. D) Extinction of conditioned freezing across 10 unreinforced CS presentations after drug treatment. E) Conditioned freezing during the test of Extinction. F) Conditioned freezing after repairing of the CS+US to test Reacquisition. Bars are means +/-SEM, each data point is a rat, n=16/group. *P<0.05

## DISCUSSION

Dropping the volume of an auditory threat cue down to just above background noise level provided a ‘vigilance screen’ that was able to parse out a range of responses in the rats to a low salience cue, some of which were as strong as if the cue was presented at full ‘normal’ volume. The overall increase in responses from the first to the second presentation (Fig 1c), suggests that any lack of initial response is not due to insensitivity to perceive the tone CS at all, but rather an attentional bias unmasks reactions to the cue presented near background noise levels that is further revealed by repeated presentations. This may be akin to an attentional tuning curve that is potentially influenced by affective state. We did not find any relation of vigilance screen behaviour to behaviour in the elevated plus maze, a task widely used to model the avoidance symptoms of anxiety disorders. We recently demonstrated a relative independence of measures from aversive conditioning and the EPM exploration (18), which supports that these tasks reveal different behavioural strategies for dealing with potential threats. Rats have been successfully studied as models for affective state biases in choice behaviours (21–23). In the experiments here, the rats in both groups were co-housed, and we can only assume under similar affective state conditions at the time of test. In future work, it would be important to ascertain whether the proportion of responses to the low salience cue can be altered by recent modulation of affective state or whether this represents a more stable phenotypic difference.

A threat cue can be rendered ambiguous because it becomes less salient or more unpredictable. Consideration of the difference sources of uncertainty has been proposed as a key refinement when looking for neural signatures related to heightened affective states (24). In this case, the outcome of the CS is likely well understood by the rats as an expectation of shock is established after conditioning, therefore the modification of the physical properties of the cue here is more likely to render it less salient rather than less predictable. The activity of the BNST was reported to be influenced by the predictability of a aversive cue (25,26). The data here adds an impact of salience on BNST activation by an aversive cue. The lateral part of the BNST negatively correlated to freezing behaviour, which may be supported by the evidence that the lateral BNST receives dense input from the infralimbic cortex, which when activated can enhance extinction of conditioned freezing (26,27).

The behaviour in the vigilance screen did not reliably predict extinction consolidation nor reacquisition. There are inconsistencies in the literature regarding whether within session extinction (acquisition) predicts subsequent extinction recall (28). Correlations between conditioning freezing at the end of extinction acquisition and the extinction test have been reported (20), but others have not found such relationships (29). Efficacy of within-session extinction is also difficult to predict in Human studies, and measurements of initial fear activation by physiological proxies were not tied to a consistent outcome (30).

Our previous work has shown that extinction persistence can be modulated by behavioural or pharmacological interventions (4). Bitopertin at the higher dose prevented alarm vocalisation, however there was not a significant effect on within-session freezing. It would be interesting to consider whether the relatively sensitivity of these readouts, freezing and USVs, reflect that some drugs may impact more the self-report of state via USVs than the implicit freezing responses. We limited our analysis to the typical 22kHz class of alarm calls in this study, though many vocalisations above this frequency were detected and recorded. Bitopertin was reported to not influence other anxiety-like readouts such as behaviour in the elevated plus maze (17). In our hands, the measurements from the EPM do not relate in a predictive matter to those from aversive pavlovian conditioning (herein, and (18)). We predicted that DCS would enhance the partial extinction of conditioning freezing, but there was no significant effect of treatment. Some rodent studies that were unsuccessful in reproducing the enhancement of extinction by DCS argued the efficacy was limited by the extent of inhibitory learning within the extinction acquisition session (31). Bitopertin on the other hand did enhance the extinction persistence at test. Both DCS and bitopertin are thought to cumulate in increased availability of the co-agonist for the glycine-binding site the GluN1-subunit containing NMDAR. NMDAR signalling is modulated by co-agonist binding (7), but the relative impact of glycine versus D-serine on subtypes of NMDAR may be critical in determining the extent of potentiation in signalling microdomains (32). Based on the good safety profile of bitopertin in Humans (14), bitopertin might be a viable candidate as a psychotherapy adjunct. Further research should investigate the impact of Bitopertin on cellular and circuit signalling in the vigilance network.

The present study aimed to investigate ultrasonic vocalisation and freezing variability under a modified aversive conditioning test to reveal neurobiology relevant to anxiety-like behaviour. These results highlight the importance of describing multiple behaviour readouts during aversive conditioning and responding, to demonstrate how threat detection is sensitive to cue salience and can be manipulated by pharmacological interventions.

## FUNDING ACKNOWLEDGMENTS AND DISCLOSURE

SD and BH are employees of and ENC has been funded by Boehringer Ingelheim Pharma GmbH & Co. KG. All authors could collegially design the study, analyse and interpret the data, write the manuscript and decide to publish the work without restriction.

## CONTRIBUTIONS

ERS and JJ performed experiments, data collection and analysis. ERS, SD and BH commented on the manuscript. ENC designed and supervised the project, performed experiments, data collection, analysis and wrote the manuscript.

## DATA AVAILABILITY STATEMENT

The datasets generated during and/or analysed during the current study are available in the Zenodo repository: https://zenodo.org/records/11093659

---

**Figure S1:**
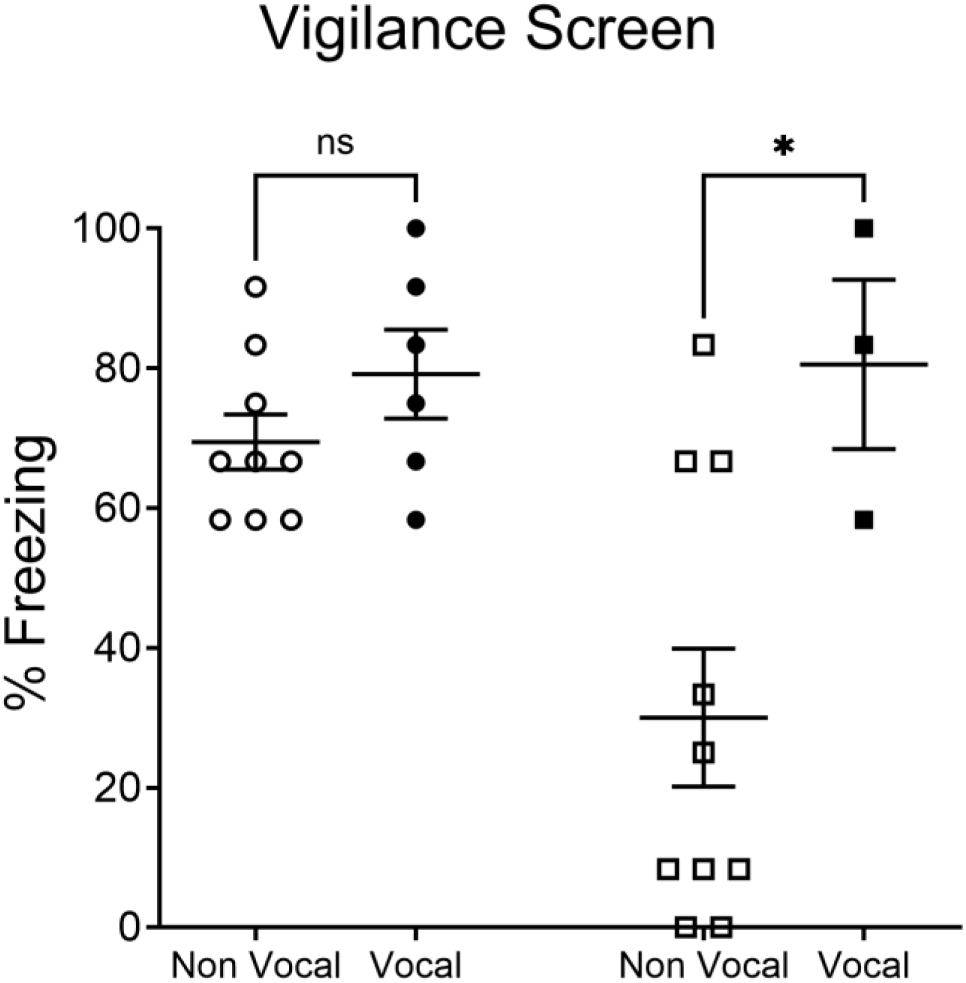
The time spent freezing by vocalising rats was significantly higher than those not emitting 22kHz alarm calls to a low salience cue. P*<0.05

